# Structure determination of a DNA crystal by MicroED

**DOI:** 10.1101/2023.04.25.538338

**Authors:** Alison Haymaker, Andrey A. Bardin, Tamir Gonen, Michael W. Martynowycz, Brent L. Nannenga

## Abstract

Microcrystal electron diffraction (MicroED) is a powerful tool for determining high-resolution structures of microcrystals from a diverse array of biomolecular, chemical, and material samples. In this study, we apply MicroED to DNA crystals, which have not been previously analyzed using this technique. We utilized the d(CGCGCG)_2_ DNA duplex as a model sample and employed cryo-FIB milling to create thin lamella for diffraction data collection. The MicroED data collection and subsequent processing resulted in a 1.10 Å resolution structure of the d(CGCGCG)_2_ DNA, demonstrating the successful application of cryo-FIB milling and MicroED to the investigation of nucleic acid crystals.

## INTRODUCTION

Determining the high-resolution structures of nucleic acids is a crucial step in understanding the structure-function relationships of these essential biomolecules. X-ray crystallography has been the primary method for studying DNA structure, RNA structure, and designed systems from DNA nanotechnology (Coimbatore Narayanan et al., 2013; Paukstelis and Seeman, 2016; Simmons et al., 2016; Subirana, 2003; Westhof, 2015). However, this method typically requires large, well-ordered crystals for collecting high-resolution data, which can pose a significant challenge for many targets. An alternative approach leverages the strong interaction of electrons with matter and their relatively lower damage to the sample (Henderson, 1995). Electron diffraction, which enables the collection of high-resolution diffraction data from much smaller crystals than those needed for conventional X-ray crystallography, is one such approach. The cryo-electron microscopy technique of microcrystal electron diffraction (MicroED) was initially developed for studying protein structures from very small microcrystals (Nannenga et al., 2014; Shi et al., 2013). Since then, MicroED has been used to solve the structures of various soluble proteins, membrane proteins, and peptides (Clabbers and Xu, 2021; Clark et al., 2021; Nannenga and Gonen, 2019).

The preparation of samples is crucial for obtaining high-resolution MicroED structures. To achieve high-quality diffraction data, it is essential to maintain the integrity of the crystal lattice while reducing the sample thickness, as the ideal thickness of a sample is only a few hundred nanometers thick (Martynowycz et al., 2021a). While there are several methods available to fragment larger crystals (de la Cruz et al., 2017) or attempt to grow small crystals, they often involve harsh treatment of the crystals or multiple rounds of sample optimization. Cryo-focused ion beam (cryo-FIB) milling has emerged as an effective technique for preparing thin lamella from biological samples as it minimizes structural damage and preserves sample hydration (Marko et al., 2007; Rigort et al., 2012). Cryo-FIB milling has been successfully used for various sample types, including protein crystals and other biological materials (Duyvesteyn et al., 2018b; Lam and Villa, 2021). The combination of cryo-FIB milling and MicroED ensures the preservation of native structures while providing high-resolution data for structure determination.

Recent studies have shown that MicroED and cryo-FIB milling are highly effective techniques for determining the structure of other biomolecular systems. For example, Martynowycz et al. (2022) reported the successful determination of the ab initio structure of triclinic lysozyme, a well-characterized model system, using MicroED at 0.87 Å resolution (Martynowycz et al., 2022). These results demonstrated the potential of MicroED to achieve atomic resolution structures from small protein crystals. Additionally, other studies such as those by Duyvesteyn et al. (2018), Martynowycz et al. (2019), Zhou et al. (2019), Polovinkin et al. (2020), and Li et al. (2018) have utilized cryo-FIB milling to prepare thin lamellae of protein microcrystals, which were then used for high-resolution structure determination by MicroED. Together with the present study, these works highlight the versatility and broad applicability of MicroED and cryo-FIB milling for determining high-resolution structures of a variety of biomolecules.

In this study, we utilized MicroED to obtain high-resolution electron diffraction data from DNA crystals, focusing on the d(CGCGCG)2 DNA duplex as our model target. This Z-DNA hexameric duplex has been extensively studied in crystallography for many years, and has been shown to diffract to very high resolution (Bancroft et al., 1994; Brzezinski et al., 2011; Chatake et al., 2005; Dauter and Adamiak, 2001; Egli et al., 1991; Gessner et al., 1989; Ohishi et al., 1991; Ohishi et al., 1996a; Ohishi et al., 2008; Ohishi et al., 2002; Ohishi et al., 1996b; Ohishi et al., 2007; Tereshko et al., 2001; Wang et al., 1979). By combining cryo-focused ion beam (cryo-FIB) milling with MicroED, we obtained a 1.10 Å resolution structure from three crystalline lamellae of this DNA duplex, demonstrating the potential of cryo-FIB milling and MicroED for nucleic acid structure determination.

## RESULTS

Crystals of the d(CGCGCG)_2_ duplex were grown using hanging drop crystallization with previously reported conditions (Brzezinski et al., 2011), yielding large crystals measuring hundreds of microns in approximately one week. Electron diffraction necessitates a sample thickness of less than a micron for beam penetration, and around a few hundred nanometers for optimal data collection (Martynowycz et al., 2021a). Cryo-FIB milling has been established as a powerful technique for thinning crystals that are too thick for MicroED analysis (Duyvesteyn et al., 2018a; Li et al., 2018; Martynowycz et al., 2019; Polovinkin et al., 2020; Zhou et al., 2019). FIB milling employs an ion beam to remove material layers without significantly damaging the unmilled sample regions. The thin crystalline lamella produced by milling can generate high-resolution electron diffraction (Martynowycz et al., 2022; Martynowycz et al., 2021a; Martynowycz et al., 2021b). Consequently, we utilized cryo-FIB milling to thin the DNA crystals for MicroED analysis.

To expedite the process of cryo-FIB milling, the large DNA crystals were initially broken into smaller fragments by pipetting them vigorously within the drops (de la Cruz et al., 2017). These fragments were promptly placed on holey carbon EM grids and vitrified using conventional MicroED sample preparation techniques (Bu and Nannenga, 2021). Upon examination of the grids in the cryo-FIB/SEM, several crystal fragments were identified on the grid. Out of these fragments, three pieces, with dimensions of approximately 30 × 15 × 10 μm, situated in the center of their respective grid squares, were selected for FIB milling, which was carried out using standard cryo-FIB milling procedures (Martynowycz and Gonen, 2021). The resulting thin lamella had a thickness of about 250 nm (Fig. 1A).

**Figure 1.**
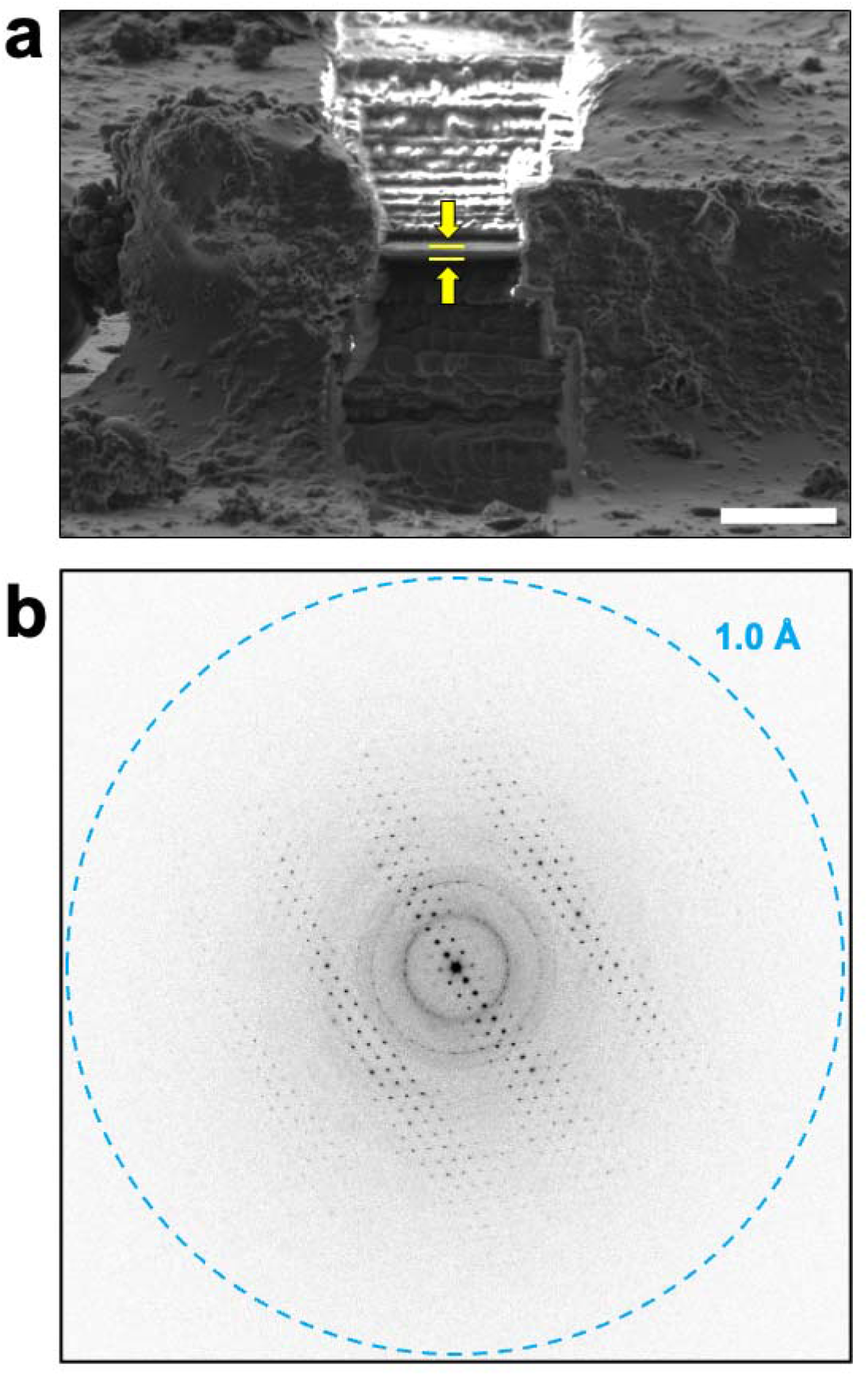
Cryo-FIB milling and MicroED data collection on d(CGCGCG)_2_ duplex crystals. a) Cryo-FIB milling was used on fragmented d(CGCGCG)_2_ crystals to prepare thin crystalline lamella of approximately 250 nm for MicroED data collection. The location of the crystalline lamella is indicated by the yellow arrows and lines, and the scale bar represents 5 μm. b) The thin DNA crystalline lamella produced by cryo-FIB milling produced high-resolution diffraction data when MicroED was performed, and continuous rotation data sets were collected from these lamellae.

FIB milled crystals were used for energy filtered MicroED data collection with a Falcon4 direct electron detector, which produced high-resolution diffraction patterns for each lamella, indicating that the diffraction power of the DNA crystals was preserved following the cryo-FIB milling process. The datasets from three lamellae were merged, resulting in a 1.10 Å resolution dataset with 98.9% completeness. The space group was *P* 2_1_ 2_1_ 2_1_ with unit cell parameters of 18.28 Å, 32.00 Å, and 41.67 Å (Table 1), in good agreement with previous studies on d(CGCGCG)2 crystals found in the PDB (Bancroft et al., 1994; Brzezinski et al., 2011; Chatake et al., 2005; Dauter and Adamiak, 2001; Egli et al., 1991; Gessner et al., 1989; Ohishi et al., 1991; Ohishi et al., 1996a; Ohishi et al., 2008; Ohishi et al., 2002; Ohishi et al., 1996b; Ohishi et al., 2007; Tereshko et al., 2001; Wang et al., 1979). Molecular replacement was used to phase the data, employing a search model of 3p4j, followed by refinement of the structure using electron scattering factors. The final structure of the d(CGCGCG)2 DNA duplex, including a spermine molecule and 33 water molecules, was determined at 1.10 Å with an overall Rwork/Rfree of 20.98 / 22.57% (Fig. 2, Table 1). Consistent with similar structures, the hexamer duplexes were found to stack along the c-axis, while the length of the a-axis corresponded to the helix’s diameter (Brzezinski et al., 2011). Notably, the structure exhibited an alternative conformation of the phosphate backbone of the third residue of chain A (Fig. X), suggesting some flexibility at this point in an otherwise rigid structure. The MicroED structure of the d(CGCGCG)2 was found to be in good agreement with previous studies on this target, and the resulting potential maps were of high quality.

**Table 1.**
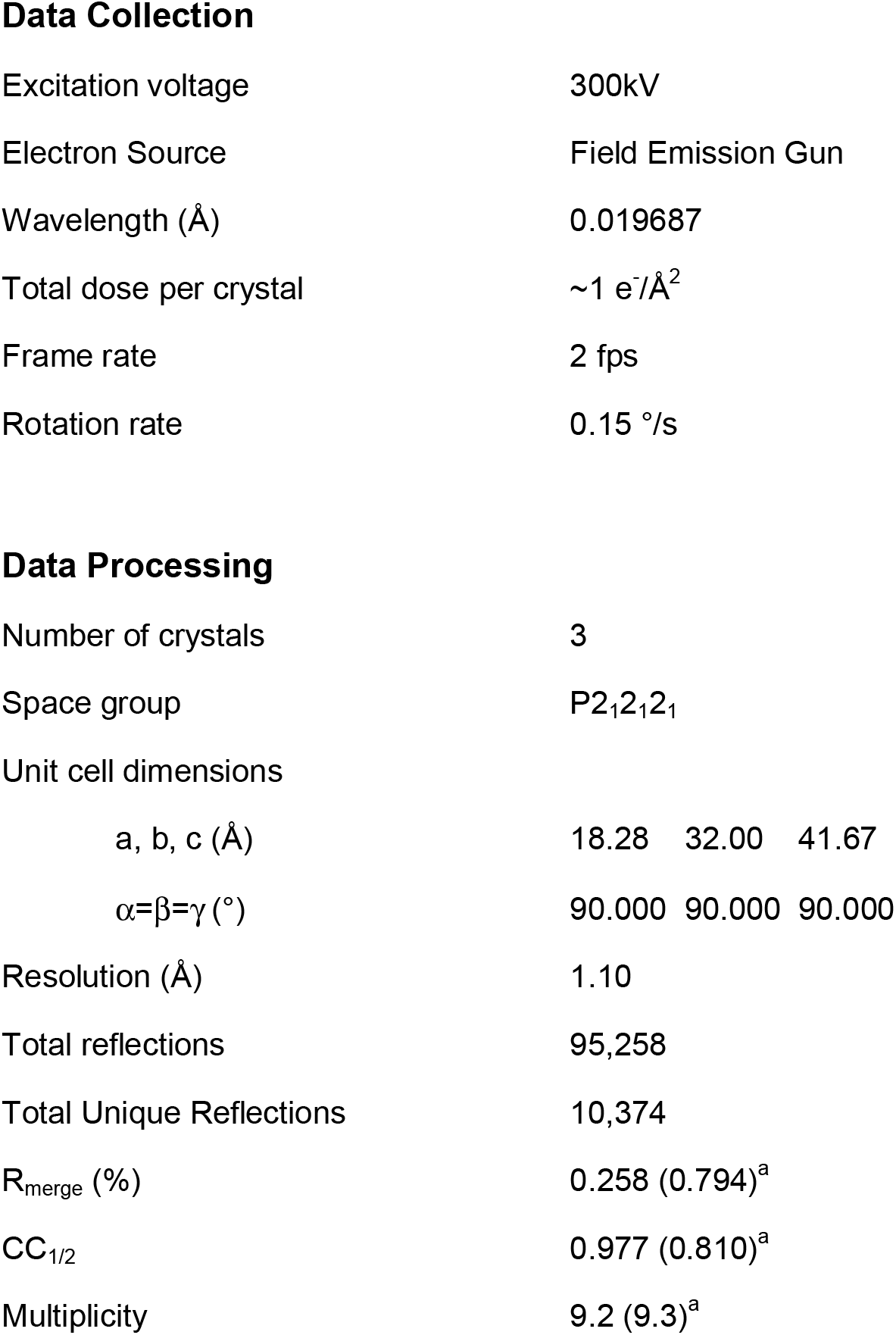
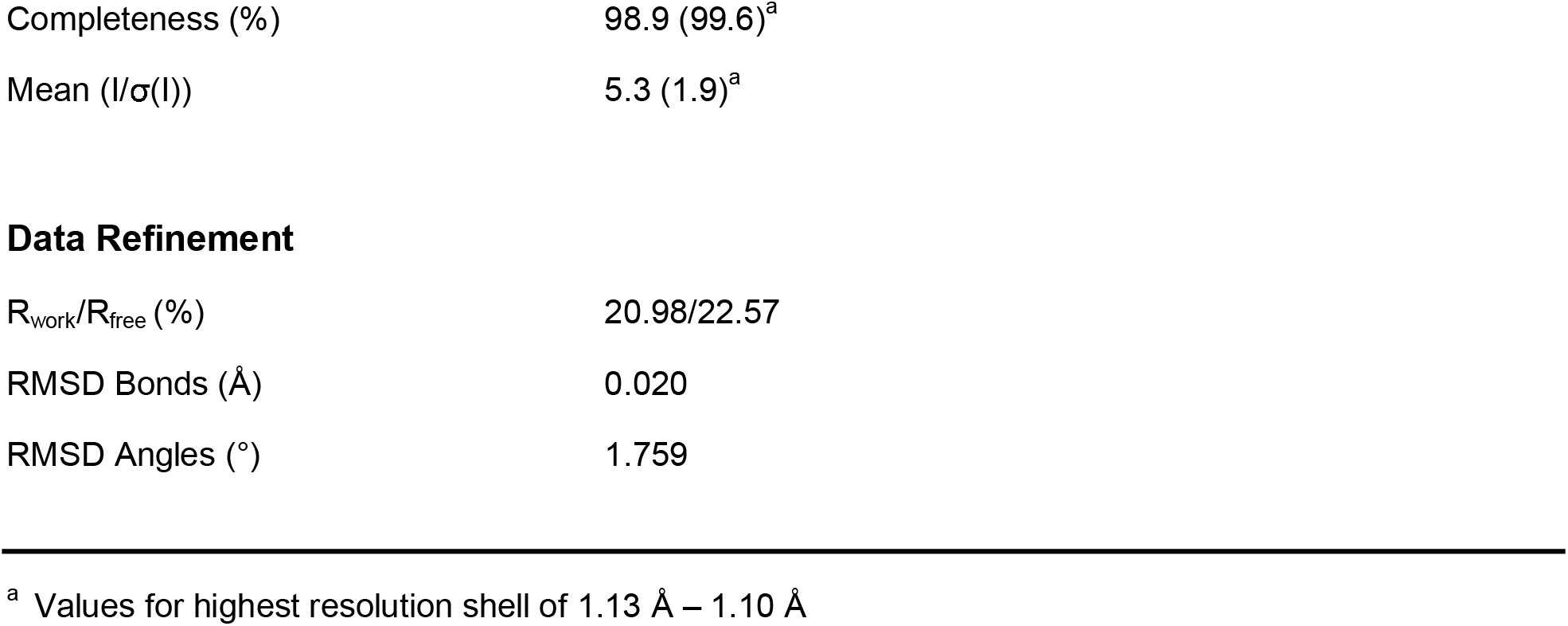
Data collection and refinement statistics.

Data Collection
|  |  |
| --- | --- |
| Excitation voltage | 300kV |
| Electron Source | Field Emission Gun |
| Wavelength (Å) | 0.019687 |
| Total dose per crystal | $\sim 1 \text{ e}^-/\text{Å}^2$ |
| Frame rate | 2 fps |
| Rotation rate | 0.15 °/s |
| Data Processing |  |
| Number of crystals | 3 |
| Space group | P2<sub>1</sub>2<sub>1</sub>2<sub>1</sub> |
| Unit cell dimensions |  |
| a, b, c (Å) | 18.28 32.00 41.67 |
| $\alpha=\beta=\gamma$ (°) | 90.000 90.000 90.000 |
| Resolution (Å) | 1.10 |
| Total reflections | 95,258 |
| Total Unique Reflections | 10,374 |
| R<sub>merge</sub> (%) | 0.258 (0.794)<sup>a</sup> |
| CC<sub>1/2</sub> | 0.977 (0.810)<sup>a</sup> |
| Multiplicity | 9.2 (9.3)<sup>a</sup> |
| Completeness (%) | 98.9 (99.6)<sup>a</sup> |
| Mean (I/$\sigma$(I)) | 5.3 (1.9)<sup>a</sup> |
| Data Refinement |  |
| R<sub>work</sub>/R<sub>free</sub> (%) | 20.98/22.57 |
| RMSD Bonds (Å) | 0.020 |
| RMSD Angles (°) | 1.759 |

<sup>a</sup> Values for highest resolution shell of 1.13 Å – 1.10 Å

**Figure 2.**
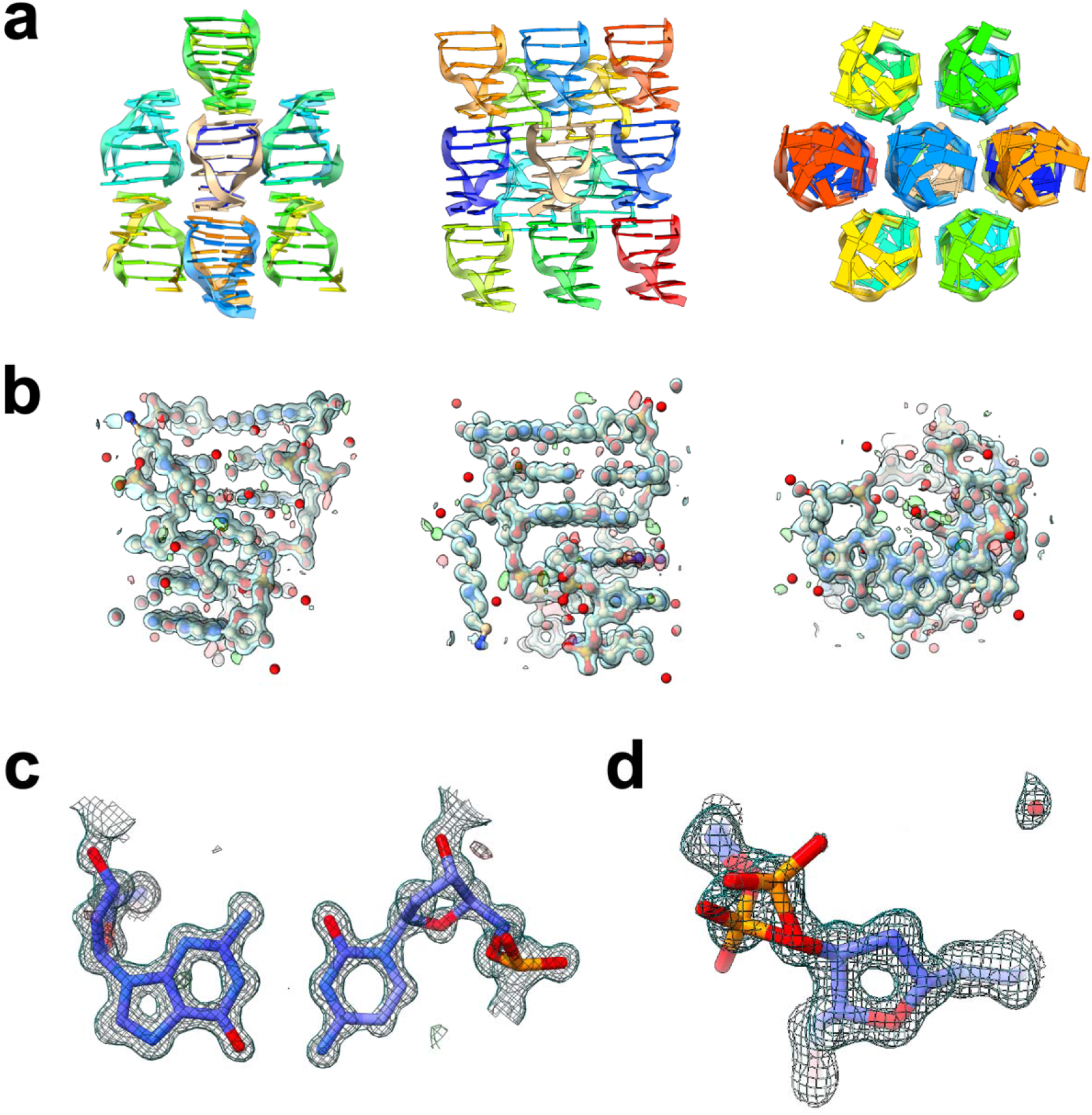
MicroED structure of the d(CGCGCG)_2_ duplex. a) The data collected from the cryo-FIB milled crystals yielded a 1.10 Å structure of the Z-DNA hexamer duplex, with packing consistent with previous studies. b) Each asymmetric unit consists of a helix of two dCGCGCG hexamers, and density is seen for each residue of the hexamers, as well as the spermine present in the structure, and several water molecules. c) the quality of the map can be clearly seen when viewing the interacting base pairs between the two hexamers. Here a representative view of residue 4 from chain A with residue 3 from chain B is shown. d) In the structure determined here, an alternative conformation of the phosphate backbone at residue 3 in chain A was modeled, where the occupancy of the two conformations were refined to values of 53% and 47%. Structures shown in (a) and (b) are viewed along the a, b, and c axes (left, middle, right, respectively). 2F_o_ – F_c_ maps shown in (b), (c), and (d) are contoured at 1.5σ (blue), and Fo – Fc maps are contoured at ± 3σ (green and red for positive and negative, respectively).

## DISCUSSION

The capability of MicroED for studying biomolecular structures from smaller crystals than those used for conventional X-ray crystallography makes the method a valuable tool for structural biology. This study highlights the applicability of MicroED to nucleic acids, which can be prepared using cryo-FIB milling. While the success of this approach in determining the structure of DNA is not surprising, it serves as a compelling demonstration of the quality of structures that can be obtained through electron diffraction data. Moreover, it expands the range of biological molecules that can be studied using MicroED, which has significant implications for future research in this field.

Compared to X-ray diffraction, electron diffraction is highly sensitive to charge effects (Wu and Spence, 2003; Yonekura et al., 2015; Yonekura and Maki-Yonekura, 2016; Yonekura et al., 2018), and nucleic acids, with their highly charged phosphate backbone, are especially affected. Nevertheless, this study demonstrates that, even with neutral electron scattering factors, the potential maps and R-factors of nucleic acids are comparable to those of protein and peptide structures determined by MicroED. However, the potential for improving the quality of nucleic acid structures using charged electron scattering factors is a potential avenue for future research as more nucleic acid structures are determined by MicroED.

The successful determination of the structure of the d(CGCGCG)2 DNA duplex using MicroED in this study suggests that MicroED can be a valuable tool for the structural analysis of nucleic acids. Further investigations on other DNA and RNA samples using MicroED could potentially yield important insights into the structural characteristics of these biomolecules. Additionally, improvements in sample preparation, data acquisition, and data processing techniques may enhance the resolution and quality of nucleic acid structures obtained using MicroED. Overall, this study highlights the potential of MicroED for nucleic acid research and encourages further exploration in this area.

## METHOD DETAILS

### DNA crystallization and grid preparation

Crystals of the Z-DNA duplex were grown following previously reported procedures (Brzezinski et al., 2011). dCGCGCG DNA was synthesized by Genscript, and a 1.5 mM solution of DNA in water was annealed at a temperature of 65 °C for 12 minutes. Following annealing, several hanging drop crystallization experiments were set up using identical conditions. 3 μL of annealed DNA was mixed with 3 μL of precipitant solution that consisted of 80 mM NaCl, 10% 2-methyl-2,4-pentanediol (MPD), 12 mM spermine tetrachloride, and 40 mM sodium cacodylate, pH 7.0. Each drop was equilibrated over 1 mL of 35% MPD at room temperature. Crystals began to appear after a few days and grew to full size in 1 to 2 weeks. For MicroED sample preparation, crystals from two drops were combined, and an equal volume reservoir solution was added to the combined drops. Crystals were fragmented by vigorous pipetting as they were monitored using a light microscope until most of the large crystals were broken down into smaller fragments. The solution containing the fragmented crystals (3 μL) was applied to the carbon side of glow discharged holey carbon grid (quantifoil 2/4, 200 mesh). Using a Leica GP2 plunge freezer, the grids were each blotted for 10 to 20 seconds prior to plunge freezing in liquid ethane.

### Cryo-FIB milling

Cryo-FIB milling was performed using an Aquilos dual beam FIB/SEM (Thermo Fisher), operating at liquid nitrogen temperatures using standard procedures for microcrystals (Martynowycz and Gonen, 2021). Briefly, vitrified grids were cryo transferred into the instrument. Whole-grid montaging using the scanning electron beam was conducted to locate crystals near the center of the grid and not near a grid bar. Each target crystal was identified using the electron beam and brought to the eucentric position. Milling was conducted at 18d in steps down to a final thickness of 2 – 300 nm. The initial, rough milling, or trenching, was conducted using a 1 nA beam current to create initial lamellae of 10um thickness. Intermediate milling was conducted by lowering the beam current as the thinned lamellae became thinner, with intermediate currents of 0.3 nA used until 3 μm, 0.1 nA until 1 μm. The 1 μm lamellae were then polished to final thicknesses using a 50 pA beam. All milling was conducted at an acceleration voltage of 30 kV, and all SEM images were acquired using a beam current of 6.1 pA and 5 kV acceleration voltage. Grids were stored in liquid nitrogen prior to future experiments.

### MicroED data collection

Following cryo-FIB milling, the grid containing the crystalline lamellae was loaded into a Titan Krios equipped with a Falcon4 direct electron detector behind a selectris energy filter operating at an acceleration voltage of 300 kV. The location of the lamella were identified at low magnification, and following alignment and setting the eucentric height, continuous rotation MicroED data were collected from each lamella (Nannenga et al., 2014). The data were collected with a rotation rate of 0.15 °/s and a camera frame rate of 0.5 frames per second, which lead to each frame in the data set spanning 0.075°. Each dataset was collected at a nominal camera length of 1200 mm, and all images were saved in MRC format and binned by 2.

### Data processing and structure determination

MicroED data collected from each of the 3 DNA crystals were indexed, integrated, scaled and merged using XDS (Kabsch, 2010). Phaser (McCoy et al., 2007) was used for molecular replacement with PDB ID: 3p4j (Brzezinski et al., 2011) used as a search model. The structure was refined using electron scattering factors with phenix.refine in the phenix crystallographic suite (Adams et al., 2010; Afonine et al., 2012) along with manual refinement using coot (Emsley et al., 2010). Structures were visualized and figures generated using ChimeraX (Pettersen et al., 2021).

## ACKNOWLEDGMENTS

This study was supported by the National Institutes of Health P41GM136508 and R01GM124152, and the Department of Defense grant MCDC-2202-002. The Gonen laboratory is supported by funds from the Howard Hughes Medical Institute.

## AUTHOR CONTRIBUTIONS

A.H., M.W.M., and B.L.N designed the experiments and protocols. A.H. and B.L.N. grew crystals and prepared samples for data collection. A.H., A.A.B, T.G., M.W.M., and B.L.N. participated in data collection, data analysis, and structure determination. All authors participated in preparing the manuscript, and all authors read and approved the final manuscript.

